# Formation of vimentin biomolecular condensate-like structures under oxidative stress

**DOI:** 10.1101/2024.05.22.595408

**Authors:** Paula Martínez-Cenalmor, Alma E. Martínez, Diego Moneo-Corcuera, Patricia González-Jiménez, Dolores Pérez-Sala

## Abstract

The intermediate filament protein vimentin performs a key role in cytoskeletal interplay and dynamics, and in cellular responses to stress. The vimentin monomer possesses a central α-helical rod domain flanked by N- and C-terminal low complexity domains. Interactions between this type of domains play an important function in the formation of phase-separated biomolecular condensates, which in turn are critical for the organization of cellular components. Vimentin filaments undergo distinct and versatile reorganizations in response to diverse stimuli. Here we show that certain oxidants and electrophiles, including hydrogen peroxide and diamide, elicit the remodeling of vimentin filaments into small particles. Diamide in particular, induces a fast conversion of filaments into circular, motile dots, for which the presence of the single vimentin cysteine residue, C328, is critical. This effect is reversible, and filament reassembly can be noticed within minutes of removal of the oxidant. Diamide-elicited structures can recover fluorescence after photobleaching. Moreover, fusion of cells expressing differentially tagged vimentin allows the detection of dots positive for both tags, suggesting that vimentin dots can merge upon cell fusion. The aliphatic alcohol 1,6-hexanediol, known to alter interactions between low complexity domains, readily dissolves diamide-elicited vimentin dots at low concentrations, whereas at high concentrations it disrupts vimentin filaments. Taken together, these results indicate that vimentin oxidation can promote a fast and reversible filament remodeling into biomolecular condensate-like structures. Moreover, we hypothesize that this reorganization into droplet-like structures could play a protective role against irreversible damage by oxidative stress.

## Introduction

Intermediate filaments constitute essential components of the cytoskeleton, from bacteria to humans (1–3). In humans, the proteins forming these fibrous polymeric structures are the products of more than sixty genes that are grouped in several classes, according to their properties (3–5). Intermediate filaments are integrators of cellular structures through their interactions with a plethora of proteins, as well as with organelles, and play key roles in mechanosensing, regulation of signal transduction, gene expression, and response to stress (reviewed in (6–8)). Certain intermediate filaments are expressed in a cell type specific manner or display a particular subcellular distribution, occurring mostly in the cytoplasm, such as the keratins, or in the nucleus, as the nuclear lamins. Notably, certain cytoplasmic intermediate filaments, such as vimentin and glial fibrillary acidic protein (GFAP), have also been found in extracellular structures, where they can participate in intercellular communication or interaction with pathogens (9–12). Vimentin belongs to the type III intermediate filament class, together with desmin, peripherin and GFAP (13). Monomers of these proteins share the basic organization of intermediate filament proteins, including a central rod segment of predominantly α-helical structure, flanked by two low complexity domains, the N-terminal, or head domain, and the C-terminal or tail domain (14). Intermediate filament assembly is a complex process that is considered to involve lateral association of parallel dimers into staggered antiparallel tetramers, several of which can further interact laterally forming unit length filaments (ULF), that may elongate by end to end annealing to constitute the filaments (3, 14).

Vimentin filaments are highly dynamic structures that undergo fast remodeling during the cell cycle as well as in response to multiple stimuli and types of stress (15–18). Vimentin normally appears in regular long filaments, extending from the periphery of the nucleus towards the plasma membrane. Nevertheless, vimentin can also reorganize in filament bundles, parallel filament arrays, juxtanuclear aggregates, cage-like structures as in aggresomes, squiggles, small particles, dots, and partly or totally diffuse structures, in response to oxidative, electrophilic, proteostatic, or osmotic stress (16, 19–21). Vimentin remodeling is finely tuned by a vast array of posttranslational modifications, which act in a concerted manner. Among them, vimentin can be phosphorylated at multiple sites, and phosphorylation has been shown to play a major role in vimentin disassembly, for instance during mitosis in certain cell types (22, 23). Importantly, vimentin is highly sensitive to redox regulation and oxidative stress. Vimentin possesses a single, conserved cysteine residue (C328) that is a key target for oxidative or electrophilic modifications that impact vimentin remodeling into morphologically distinct patterns (18, 20, 21, 24). In particular, certain electrophiles elicit the retraction of vimentin filaments towards the nucleus, whereas others can induce bundling or filament dissociation into dots (21, 25, 26). Given the interactions of vimentin with numerous cellular proteins and organelles, its remodeling likely affects the subcellular distribution of cellular components, as it has been evidenced during aggresome formation, or upon heat shock elicited filament retraction, which can provoke the concentration of associated proteins such as chaperones (8, 20, 27, 28).

The organization of cellular material is critical for cellular functions. Membraneless organelles are receiving increased attention as a means to generate intracellular compartments with important functions in stress responses, regulation of gene transcription or genome organization, as well as in the mechanisms of disease (29, 30). The process known as phase separation or biomolecular condensation, is crucial for the generation of these membraneless compartments. Importantly, the centrosome, nucleolus and stress granules are examples of biomolecular condensates (31, 32). The capacity of protein domains of low sequence complexity to self-associate through labile interactions is frequently at the basis of phase separation. Interestingly, studies with the isolated head domains of vimentin, peripherin and other intermediate filaments have shown that they phase separate forming gel-like condensates, which has been reported to involve the formation of labile cross-ꞵ structures (33, 34). Moreover, labile interactions of the low complexity head domain of intermediate filaments have been proposed to assist in filament assembly (34, 35). This conclusion is also supported by the recently available model of the vimentin filament structure, obtained by cryo-EM tomography, according to which the head domains are projected towards the center of the filament where they associate (36). Taken together, these observations support an important role of the self-association properties of the head low complexity domains in the dynamic regulation of vimentin and other intermediate filaments (35).

The fast remodeling of vimentin in response to oxidative stress or to certain electrophiles constitutes an excellent example of the dynamicity and versatility of the vimentin network (18, 37, 38). Particularly intriguing is the capacity of vimentin filaments to disassemble into dots or droplet-like small particles in response to certain oxidants (20, 21, 26). Here, we have addressed the hypothesis that these structures could behave as condensates. Our results indicate that oxidant-elicited vimentin dots share characteristics of biomolecular condensates, including their apparently spherical shape, ability to fuse and exchange material with the surroundings. Moreover, these dots are reversible structures, indicating that vimentin assembly can be recovered. Taken together, these observations suggest that vimentin in condensate-like dots could constitute a reservoir for filament recovery after oxidative stress.

## Materials and methods

### Reagents

Diamide, monobromobimane, 1,6-hexanediol and 2,5-hexanediol were from Sigma. H_2_O_2_ and anti-vimentin V9-Alexa 488 (sc-6260) were from Santa Cruz Biotechnology.

### Cell culture and treatments

DMEM, penicillin/streptomycin and trypsin/EDTA were from Gibco. Fetal bovine serum (FBS) was from Sigma. Cells were cultured at 37°C in a humidified atmosphere with 5% CO_2_. SW13/cl.2 vimentin deficient cells were the generous gift from Prof. A. Sarriá (University of Zaragoza, Spain). SW13/cl.2 cells stably expressing vimentin wild type (wt) or vimentin C328A have been reported previously (20, 21).

Cells were cultured in DMEM supplemented with 100 U/ml penicillin and 100 μg/ml streptomycin, and 10% (v/v) FBS. Treatments were performed in the absence of serum. Diamide was used at 1 mM, final concentration. When specified, medium containing diamide was exchanged with fresh serum-free medium. Aliphatic alcohols were used at 3.3% to 8% (w/v), final concentrations, by dilution from a 50% (w/v) stock solution prepared in water or in serum-free culture medium.

### Plasmids and transfections

The fusion plasmid GFP-vimentin wt and the bicistronic plasmids coding for DsRed-Express2 and vimentin wt or C328A mutant (RFP//vimentin wt or C328A), have been described previously (20, 21). The RFP//vimentin L387P mutant was generated by site-directed mutagenesis using oligonucleotides 5’-CGT GAA TAC CAA GAC CTG CCC AAT GTT AAG ATG GC-3’ and its complementary reverse, and the NZYMutagenesis kit from NZYtech, following the instructions of the manufacturer. mCherry-vimentin was from Genecopoeia. Amplified plasmids were purified using the NZYMaxiprep Endotoxin free kit from NZYtech. SW13/cl.2 cells were transfected using Lipofectamine 2000 (Invitrogen), using a proportion of 3 µl per microgram of DNA, both prepared in Optimem, following the manufacturer instructions. For live cell monitorization of vimentin, cells were transfected with a proportion of 80% untagged vimentin plasmid (RFP//vimentin) and 20% GFP-vimentin, as previously reported (18). Cells were incubated with the transfection mixture for five hours, after which, the medium was replaced with fresh medium and incubation was continued for 48 h in the absence of antibiotics. Transiently transfected cells were routinely used 48 h after transfection.

### Fusion assays

For cell fusion, a modification of a previously published procedure was used (39). Briefly, on day 1, cells were transfected with RFP//vim wt (0.8 µg) + GFP-vim wt (0.2 µg) or with RFP//vim wt (0.8 µg) + mCherry-vim wt (0.2 µg), as above, to achieve the formation of GFP or mCherry labeled vimentin networks, respectively. On day 2, cells expressing GFP or mCherry labeled vimentin were trypsinized and plated together in a p35 dish to achieve a mixed population. On day 3, cells were treated with 1 mM diamide for 15 min to elicit the formation of vimentin dots. Immediately afterwards, cell fusion was performed by incubating the dish with 1 ml of prewarmed polyethylene glycol 1500 (PEG 1500, Roche) for 1 min at 37°C. PEG was removed carefully and after 1 min, medium containing diamide was added and incubation continued at 37°C for the indicated times. Monitorization of fusion was performed in live or fixed cells, as specified.

### Immunofluorescence and microscopy

Cells cultured on glass coverslips (Epredia Netherlands B.V., Ø12 mm, #1.5) were treated with the various agents, washed twice with PBS and fixed with 4% (w/v) paraformaldehyde (PFA) for 25 min at room temperature. Immunofluorescence was carried out essentially as described (21). Briefly, cells were permeabilized with 0.1% (v/v) Triton X-100, blocked with 1% (w/v) bovine serum albumin (BSA) in PBS and incubated with anti-vimentin Alexa 488 at 1:200 dilution in blocking buffer. Coverslips washed with PBS and water were allowed to dry and mounted on glass slides with FluorSave^TM^ Reagent (Merck). Cells were visualized on Leica SP5 or SP8 confocal microscopes. Routinely, single sections were acquired every 0.5 µm. Single confocal sections or overall projections are shown as indicated.

For FRAP assays, cells were seeded on glass bottom cell culture dishes (Mattek Corporation) and transfected with RFP//vimentin plus GFP-vimentin, as detailed above. Cells were treated with diamide 1 mM, final concentration, for 15 min, and assays were performed immediately afterwards in a Leica SP5 microscope equipped with a thermostatized support. Briefly, after acquiring a pre-bleach image with a zoom of 4x, a region of interest (ROI) of 10 x 2.8 µm was bleached by three “shots” of the green laser of a Leica SP5 microscope at 75% potency. A postbleach image was immediately obtained and then single sections were acquired every five seconds during three minutes. Fluorescence intensity of the ROI from these assays was quantitated with LasX software.

Image analysis was performed with Leica Las X or Image J (FIJI) software. The Analyze Particles menu of Image J was used for calculation of particle size and circularity. Particle tracking was performed with the Trackmate or the Manual tracking plugins.

### Statistical analysis

All experiments were repeated at least three times and representative results are shown. Statistical analysis was performed with GraphPad 9.0 software. Comparisons between two data sets were performed by the Student’s *t*-test for unpaired samples, followed by Kolmogorov-Smirnof test. Statistically significant differences are indicated on the graphs and/or in the figure legends, as follows: *p<0.05, **p<0.01, ***p<0.001, ****p<0.0001. The number of determinations or the sample size are given in the figure legends.

## Results

### Various cysteine-reactive agents elicit partial vimentin remodeling into small particles

We have previously reported that treatment with various oxidative or electrophilic compounds elicits profound and characteristic remodeling of the cellular vimentin network (21). In several instances, the remodeling pattern is heterogeneous and consists of morphologically diverse elements, including bundles, diffuse material, linear arrays, or dots and aggregates of variable size. Here, we observed that treatment of SW13/cl.2 cells stably expressing vimentin wt (20) with various oxidants or cysteine-reactive agents, such as the reactive oxygen species H_2_O_2_ or the cysteine reagent monobromobimane (MBrB), elicits filament bundling, juxtanuclear accumulations, and/or retraction from the cell periphery (Fig. 1A). Notably, in cells undergoing filament retraction, the peripheral area was devoid of full filaments when visualized under non-saturation conditions for the whole cell. However, increasing the exposure of the peripheral area allowed the detection of numerous vimentin small particles of variable size, indicative of filament disassembly or disengagement (Fig. 1A). Moreover, aligned particles indicative of filament remnants were also apparent (Fig. 1A, insets), whereas these strings of dots were mostly absent in control cells. Remarkably, and consistent with our previous observations (20), the oxidant diamide-elicited an extensive reorganization of the vimentin network into small, nearly spherical particles or dots (Fig. 1A, right panel), which appear distributed throughout the cytoplasm. Given the robustness of the effect of diamide, we studied this remodeling process in more detail. Image analysis of diamide-elicited vimentin dots showed that more than 95% of the particles displayed an area <0.5 µm^2^, with an average size of 0.116 ± 0.001 µm^2^. Interestingly, a considerable proportion of the dots (nearly 30%) displayed a size lower than 0.05 µm^2^, which is compatible with the size of one or a few ULF. In fact, ULF size in cells appears to display a heterogenous distribution with estimated average lengths ranging from ∼59 to 130 nm, depending on the experimental setting (40, 41). These estimations would imply an area lower than 0.017 µm^2^. Moreover, dots <0.5 µm^2^ exhibited a consistent circular shape (average circularity of 0.86 ± 0.002), with 47.3 % of them displaying a value of 1 (complete circularity). In contrast, the larger structures remaining after a 15 min diamide treatment had a poor circularity (0.54 ± 0.009) and a morphology characteristic of incomplete filament dissociation or aggregation of several dots.

**Figure 1.**
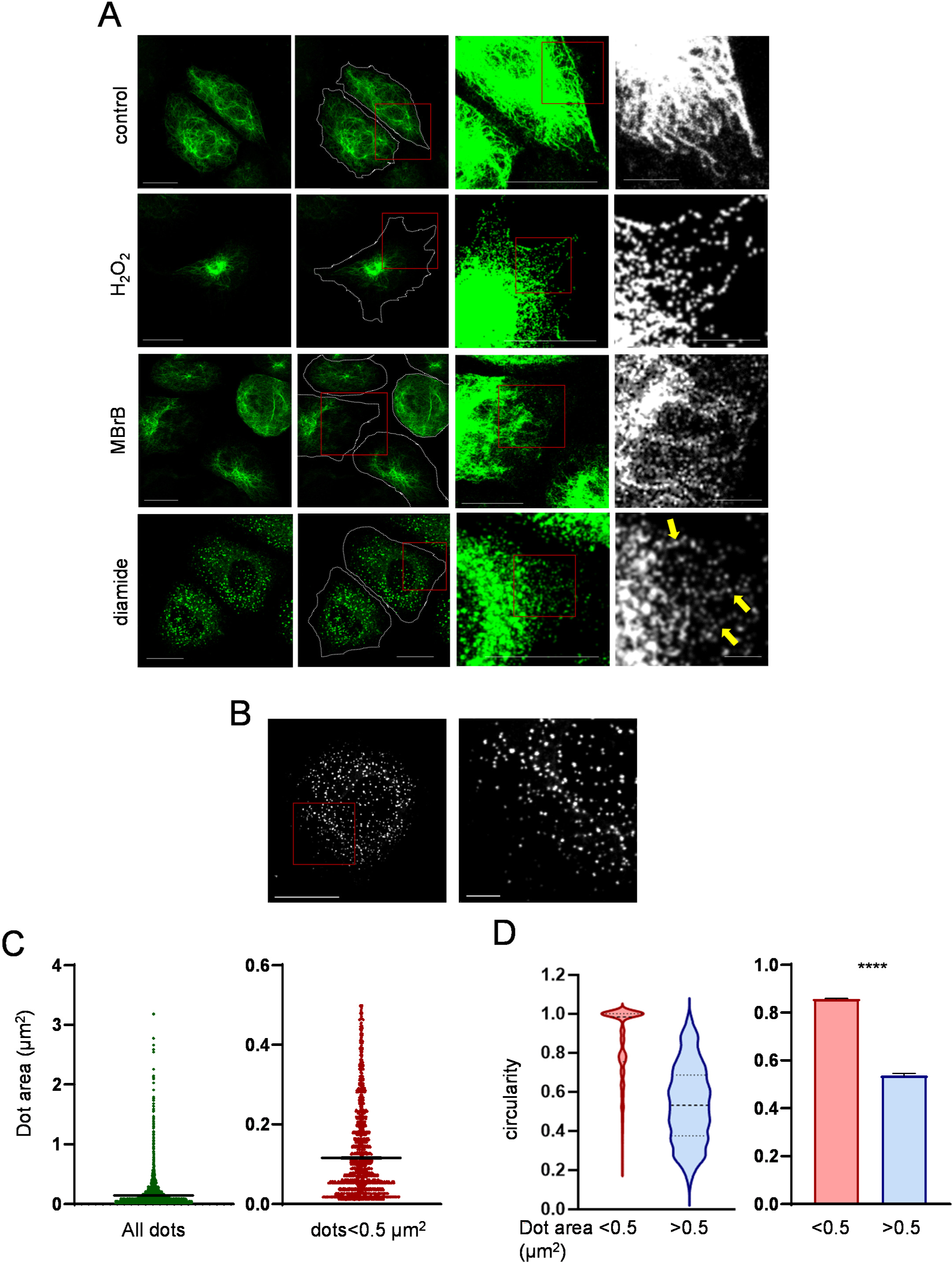
Various oxidants elicit vimentin filament remodeling into dots. (A) SW13/cl.2 stably transfected with untagged vimentin (RFP//vimentin wt plasmid), were treated with 1 mM H_2_O_2_ for 30 min, or 1 mM MBrB or 1 mM diamide for 15 min. Vimentin structures were visualized by confocal fluorescence microscopy after immunofluorescence. Images shown are overall projections. From left to right, raw images of cells; images with the cell contour outlined and a region of interest (ROI 1) delimited; enlarged images of ROI 1 deliberately overexposed to show the vimentin particles at the peripheral region of cells, and with a second ROI (ROI 2) delimited; detail of ROI 2. Arrows mark filament remnants appearing as strings of dots. Scale bars, 20 µm in the three left columns and 5 µm in the right column. (B) Vimentin dots in a cell treated with diamide for 15 min. The inset is enlarged at right. Scale bars, 20 µm (left image) and 5 µm (right image). (C) and (D) Characterization of vimentin dots. (C) The area of dots present in 20 cells from 3 experiments, totaling 11413 dots, is depicted at the left. The graph at the right provides a more detailed view of the area of dots smaller than 0.5 µm^2^, which account for 95% of total dots. (D) The circularity of vimentin dots depending on their size (smaller or larger than 0.5 µm^2^) is shown as violin plots of the two populations (left graph), and as average values ± SEM from 10906 and 507 values, respectively. ****p<0.0001 by unpaired Student’s t-test followed by the Kolmogorov-Smirnof test.

### The reorganization of vimentin filaments into dots elicited by diamide is reversible

Diamide is an electrophilic compound that elicits the formation of disulfide bonds (18, 42, 43). This is actually the main modification of vimentin observed upon treatment with diamide in vitro (18, 26). This modification is readily reversed under reducing conditions. Therefore, we assessed the potential reversibility of diamide-elicited vimentin remodeling by monitoring vimentin organization in cells treated with diamide for various intervals, as well as after diamide removal, as schematized in Fig. 2A. Diamide showed a time-dependent effect, which progressed from partial fragmentation of vimentin filaments after a 5 min treatment, to their nearly complete conversion into dots after 15 min, and the subsequent appearance of a diffuse cytoplasmic vimentin background, detectable at 30 min, which further increased after 2 h treatment (Fig. 2B, upper images). Examination of the vimentin network at several time points after diamide removal showed that vimentin dots could reconstitute long filaments. Recovery was more efficient after short diamide treatment. Indeed, vimentin dots elicited by a 5 min treatment with diamide turned into a network of connected squiggles or short filaments 10 min after removal of the oxidant, they later formed wavy filaments, and finally, long robust filaments at the end of a 115 min recovery period (Fig. 2B, middle images). Conversely, when recovery was started after a 15 min diamide treatment, vimentin dots initially transformed into larger irregular accumulations (15 min recovery), which were later connected by numerous filamentous structures (115 min recovery, Fig. 2B, lower images). The appearance of filaments after diamide removal indicates that vimentin present in dots may still be functional and not severely damaged. Nevertheless, the less efficient recovery of the network after longer treatment with diamide may indicate that additional modifications and/or disruption of other cellular structures or signaling pathways had occurred.

**Figure 2.**
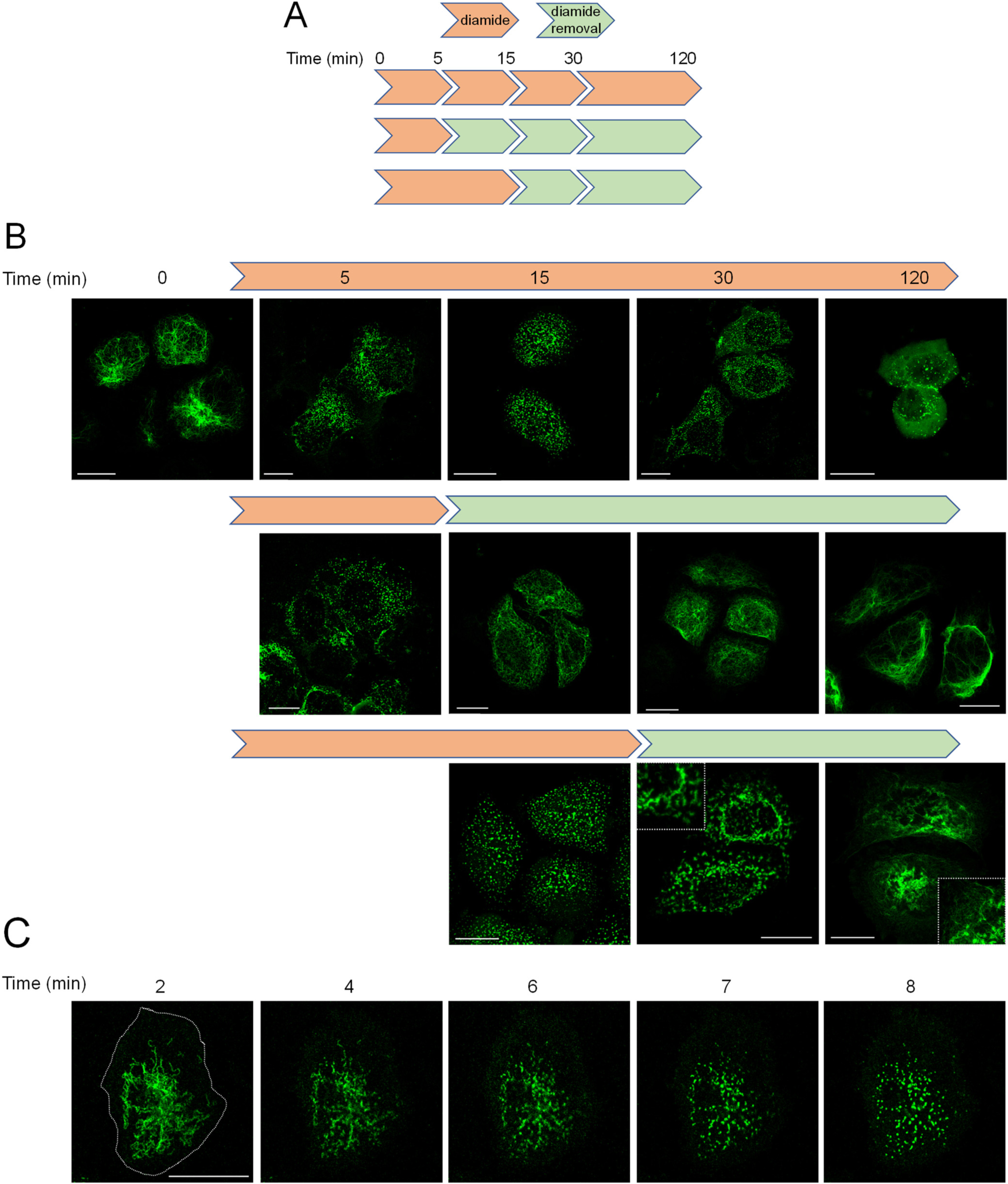
Reversibility of the effect of diamide. (A) Scheme of the experimental design displaying the times for incubation in the presence and in the absence of diamide. (B) SW13/cl.2 cells stably expressing vimentin were treated with diamide for the indicated times. At the times specified in the scheme in (A), diamide was withdrawn and cells allowed to recover in serum-free medium. At the end of the 2 h total incubation, cells were fixed and vimentin distribution was assessed by immunofluorescence and confocal microscopy. Overall projections are shown. Images are representative from at least three experiments with similar results. (C) Monitorization of vimentin filaments dissociation in live cells. Cells were transfected with a combination of untagged and GFP-tagged vimentin plasmids as detailed in Methods, and the distribution of vimentin was monitored during treatment with diamide. Images (single sections) were acquired at the indicated time points. Bars, 20 µm.

Fixation procedures can fail to provide a fast snapshot of biomolecular interactions, particularly those that are transient in nature, and/or alter the appearance of transient structures (44). Therefore, we monitored diamide-elicited vimentin filament conversion into dots in live cells. For this, cells were transiently transfected with vimentin wt and a small amount of GFP-vimentin wt (18, 20). This approach allowed the direct observation of the reorganization of filaments into dots in a matter of minutes (Fig. 2C and Suppl. Video 1). This confirmed that the observations in fixed cells closely reflected the process of diamide-elicited vimentin remodeling. In view of these observations, it could be considered that, more than severing into short filaments, diamide-elicited vimentin remodeling results in a dissociation of filaments into small structures that retain the capacity to reassociate.

### Diamide-elicited vimentin dots are motile

Interestingly, observation of diamide-elicited vimentin dots in live cells showed that they are motile structures (Fig. 3). Both, oscillations and displacements, in some cases covering several micrometers, were observed. Examples of particle displacements occurring in a time frame of seconds (upper images) or minutes (lower images) are shown in Fig. 3A. Video recordings of cells showing the complete image sequences are included in the Supplementary material (Suppl. Video 2 and 3). Analysis of the movement of particles randomly selected from several experiments, depicted in Fig. 3B, indicates that dots move actively, although the majority of them remain within a radius of 1 or 2 µm from the point of origin. Nevertheless, a population of particles that end up at distant points of the cell could be clearly identified. Monitorization of the movement of particles in a single cell is illustrated in Fig. 3C. In this representation, red lines highlight trajectories of particles that have traveled a distance close to 10 µm, independently of the direction. Trajectories up to 10 µm, which account for 96% of total, are represented in Fig. 3C. The movement of particles along the highlighted trajectories can be followed in Suppl. Video 4. A distribution of the speed of the particles is shown in Fig. 3D. Taken together, these observations show that diamide-elicited vimentin dots are highly motile structures.

**Figure 3.**
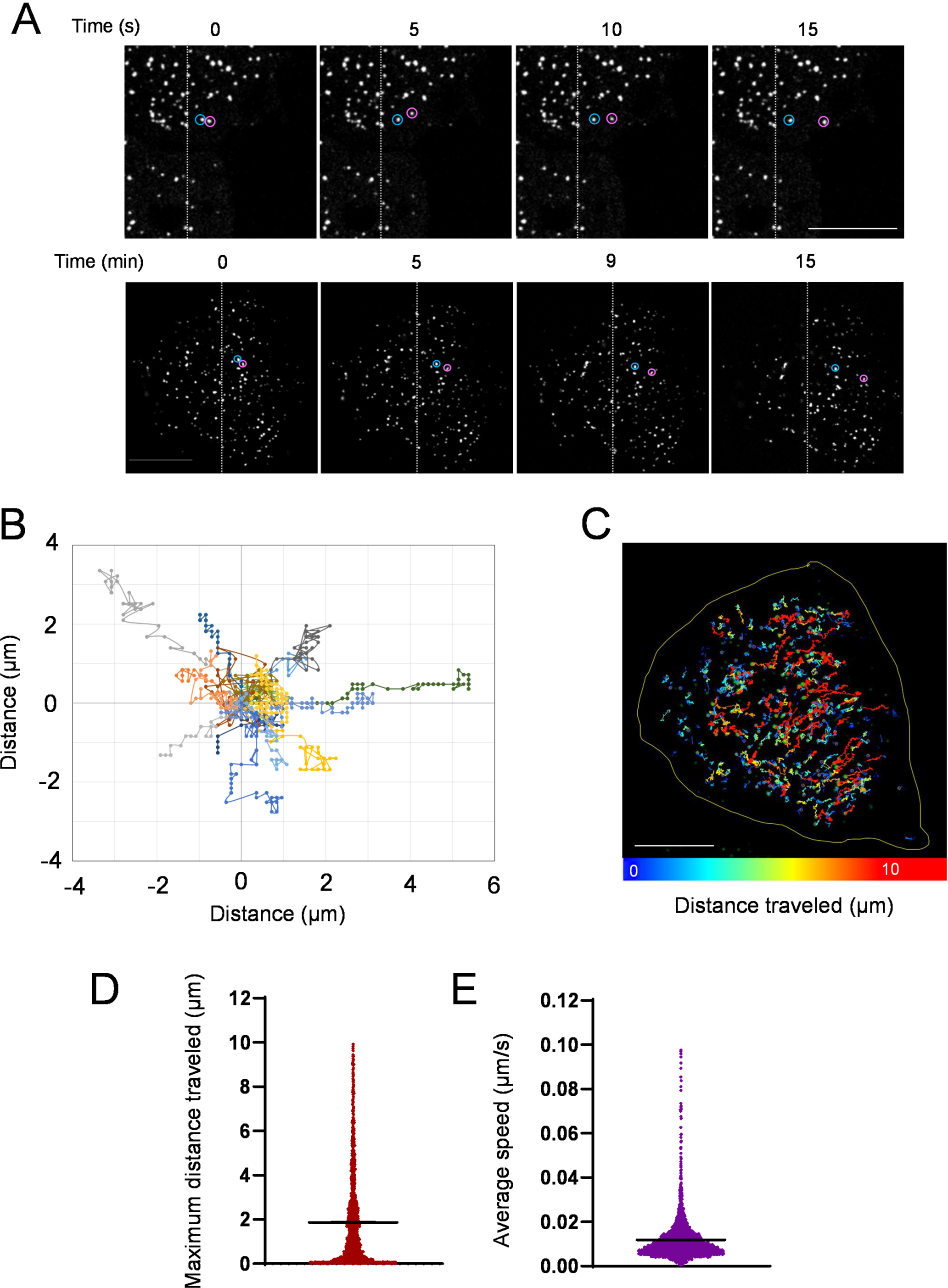
Motility of vimentin dots. Cells were transfected with a combination of plasmids coding for untagged and tagged vimentin, as detailed in Methods, and monitored live after treatment with diamide for 15 min. Images were acquired every ten seconds and selected sequences are shown to illustrate fast (upper row) and slower (lower row) dot displacements. A vertical line is drawn on images for reference. In addition, selected dots are encircled in blue (reference dot) and pink (moving dot). Bars, 10 µm. (B) Graph representing the movement of aleatorily selected dots from 17 cells from two experiments. (C) Track analysis of dots present in a single cell. The length of the trajectories for every dot monitored is represented by a color code, with trajectories of approximately 10 µm appearing in red. The pseudocolor scale is shown at the bottom. Bar, 10 µm. (D) and (E) Characterization of the motility of diamide-elicited vimentin dots. (D) Distance covered by dots is represented as average value ± SEM of 2422 determinations from three experiments (trajectories up to 10 µm are represented). (E) Average speed of dots, represented as mean value ± SEM of 2499 determinations. For clarity of the graphs, dots with average speed ≤ 0.1 µm/s, which account for 99%of particles, are represented.

### Vimentin dots are dynamic structures that undergo FRAP

Assessment of the mobility or dynamicity of components within biomolecular condensates is often assessed by fluorescence recovery after photobleaching (FRAP) (45). Therefore, we explored whether diamide-elicited vimentin droplets could undergo FRAP. FRAP experiments clearly showed recovery of fluorescence early after photobleaching, with a significant increase in fluorescence being detected at the first time point registered after bleaching (5 s) (Fig. 4). Representative pre and postbleach images are shown in Fig. 4A, whereas Fig. 4B shows a typical recovery sequence. The complete video sequence is shown in Suppl. Video 5. The graph in Fig. 5C depicts the quantification of fluorescence recovery of bleached areas from several experiments. These results indicate that there is exchange of GFP-vimentin containing “subunits” between the vimentin dots and the unbleached surrounding environment (46).

**Figure 4.**
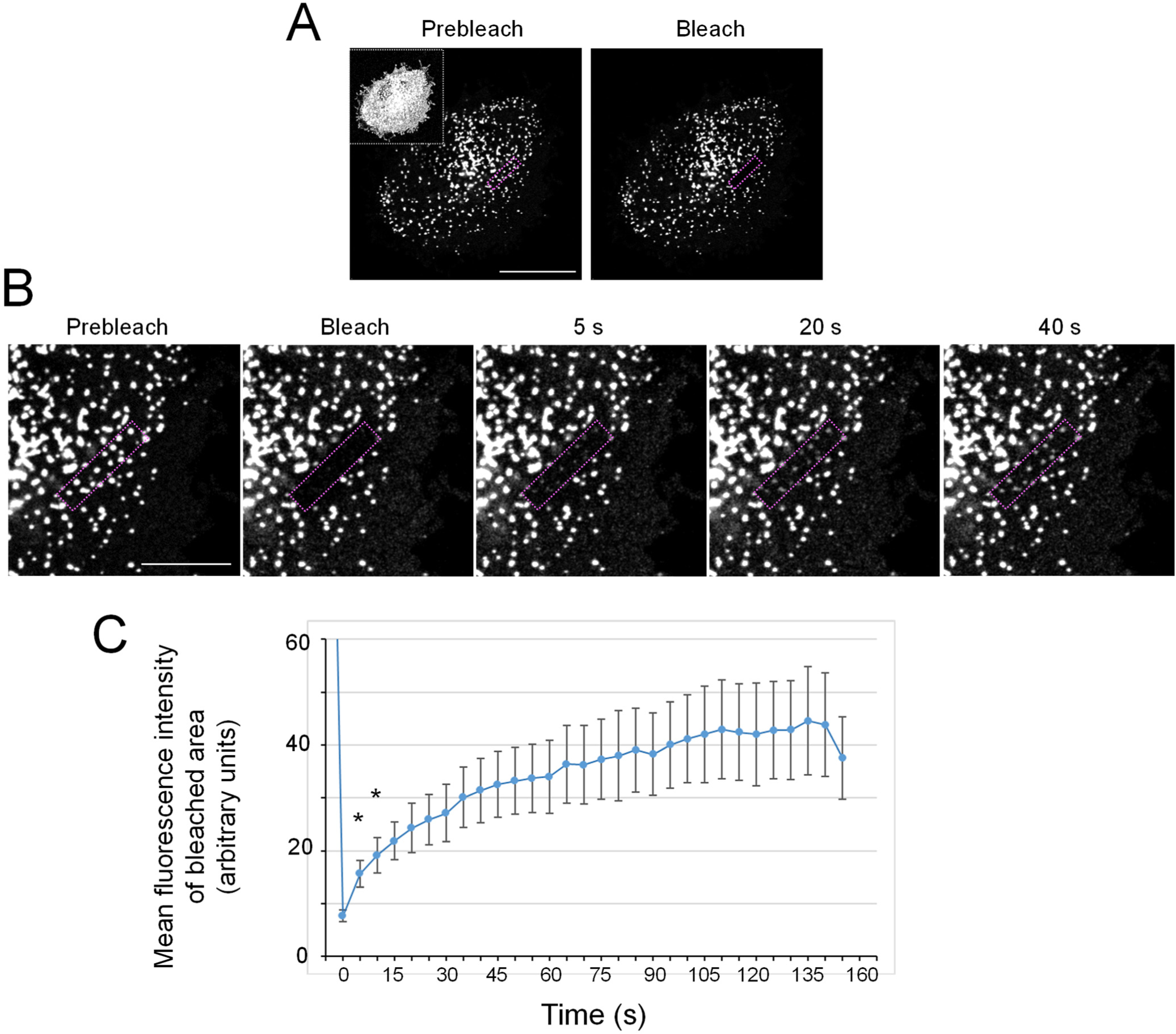
Fluorescence recovery after photobleaching (FRAP) of diamide-elicited vimentin dots. Cells expressing a combination of untagged and GFP-tagged vimentin were subjected to FRAP assays as detailed in the experimental section. (A) Prebleach and immediately postbleach acquired images are shown. The bleached area is highlighted by a dotted rectangle. Bar, 20 µm. The inset shows an overexposed image of the cell to illustrate the cell contour. (B) Sequence of enlarged images showing the detail of the bleached area. Single sections of the area of interest were acquired every 5 s and images corresponding to the indicated time points are shown. Bar, 10 µm. (C) Quantitation of fluorescence recovery. Average fluorescence intensity of the beached area at the indicated time points was quantitated using Image J, and mean values from 17 FRAP assays ± SEM are represented. *p<0.05 with respect to the postbleach (0 time) fluorescence intensity. Note that the increase in fluorescence intensity at all time points is statistically significant with respect to 0 time, although it is only specified for the 5 and 10 s time points for clarity.

**Figure 5.**
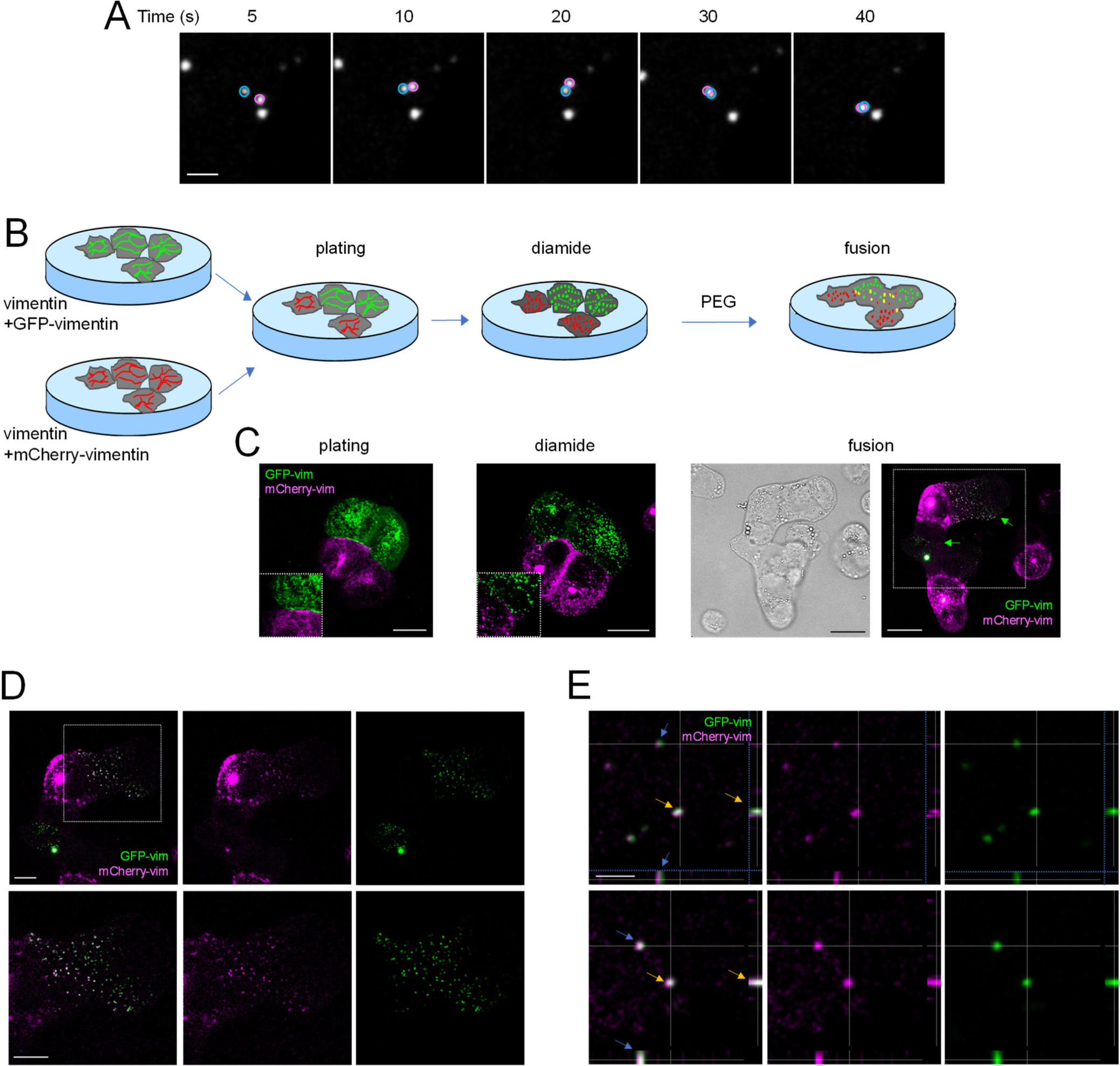
Merging of differentially tagged vimentin dots upon cell fusion. (A) Monitoring of live cells shows collision of vimentin dots. Cells transfected with a combination of untagged and GFP-tagged vimentin were monitored, and images acquired at the indicated times are shown. Selected dots are encircled in blue or pink lines to illustrate their coalescence (30 s), and subsequent joint displacement (40 s). Bar, 2 µm. (B) Scheme depicting the strategy for cell fusion of cells expressing differentially tagged vimentin. Briefly, cells transfected with combinations of untagged vimentin and GFP-or mCherry-tagged vimentin to produce green or red “colored” filaments were plated together, treated with diamide to elicit the formation of dots, and subsequently fused with PEG. (C) Representative images (overall projections) of the different steps in the fusion process. In “plating” and “diamide”, insets show enlarged areas illustrating differentially tagged filaments and dots, respectively. In “fusion”, the left image shows the bright field illustrating the continuity of the cell membrane and presence of multiple nuclei in fused cells; the right image depicts the different tagging of vimentin in the fused cells. Bars, 20 µm. Green arrows point to cells that apparently expressed GFP-tagged vimentin, originally. The area denoted by the dotted rectangle is enlarged in (D). (D) Images showing single sections of the merged and individual channels of fused cells with vimentin dots bearing GFP and mCherry tags in different proportions. Inset is enlarged at the bottom images. Bars, 10 µm. (E) Images of selected vimentin dots are shown to illustrate the different degrees of overlap between the GFP and mCherry signals. Single sections and the corresponding orthogonal projections are shown. Individual dots are identified on the projections by color-coded arrows. Bar, 2 µm.

### Fusion of adjacent cells with differentially tagged vimentin dots results in the appearance of structures bearing both tags

Observation of vimentin dots in live cells showed that some of them apparently collided and subsequently moved together (Fig. 5A). Therefore, we attempted to explore whether they could undergo fusion. For this, cells cotransfected with untagged vimentin plus GFP-or mCherry-vimentin were plated in the same dish, treated with diamide and subjected to a fusion process with polyethylene glycol, as detailed in Fig. 5B. This process elicited the appearance of fused cells, distinguishable by a continuous plasma membrane as well as the presence of several nuclei, in which cytoplasmic contents from the original cells could be exchanged. Representative images from each one of these steps, shown in Fig. 5C, illustrate that GFP and mCherry-tagged vimentin filaments behaved similarly in response to diamide. Interestingly, fusion of cells expressing differentially tagged vimentin led to the appearance of dots bearing both GFP-vimentin and mCherry-vimentin (Fig. 5C). Certain areas of fused cells showed a gradual mixing of green and red dots (Fig. 5D). Moreover, individual dots displayed a variable overlap of the green and red tags, as evidenced in the orthogonal projections (Fig. 5E). These results suggest that coalescence between the two populations of dots can occur. Nevertheless, the possibility that exchange of vimentin subunits also contributes to the appearance of mixed dots cannot be discarded.

### Diamide-elicited vimentin dots are selectively dissolved by 1,6-hexanediol

Several protein condensates have been shown to be susceptible to “dissolution” or “melting” by certain aliphatic alcohols in a structure-dependent manner. The basis for this effect appears to be the disruption of weak hydrophobic interactions (47). Of the alcohols employed in various studies, 1,6-hexanediol appears particularly effective for melting cross-ꞵ polymers formed from low complexity domains, hypothetically due to the spacing between its hydroxyl groups, whereas 2,5-hexanediol is much less effective (33). Therefore, we explored the impact of 1,6-hexanediol and 2,5-hexanediol on diamide-induced vimentin dots (Fig. 6A). Incubation of diamide treated cells with 3.3% (w/v) 1,6-hexanediol for 5 min completely dissolved vimentin dots, leading to the appearance of diffuse cytoplasmic vimentin, whereas 2,5-hexanediol had no effect. Interestingly, a previously described vimentin L387P mutant, involved in the pathogenesis of a rare premature aging syndrome, forms aggregates when expressed in vimentin deficient cells (48). As a control of the selectivity of the effect of aliphatic alcohols, we assessed their impact on the aggregates formed by vimentin L387P. We observed that these aggregates were not dissolved by treatment with either of the alcohols (Fig. 6B), which indicates that they possess different structure and/or composition than diamide-elicited vimentin wt dots. Remarkably, at 3% (w/v), neither alcohol altered vimentin wt filaments per se, even after a 20 min incubation (Fig. 6C). Nevertheless, the vimentin network has been reported to be dispersed into small puncta by exposure to higher concentrations (8% (w/v)) of 1,6-hexanediol (33), which has been interpreted as a consequence of the disruption of cross-ꞵ interactions between the N-terminal domains, which play a role in filament assembly (33). Therefore, we explored the effect of increasing concentrations of these alcohols in our experimental model. Here we observed that increasing the 1,6-hexanediol concentration to 5% resulted in a mixed vimentin pattern, ranging from preservation of filaments (Fig. 6D, blue arrow) to reorganization into small fragments or particles, and even appearance of cells with diffuse vimentin (Fig. 6D, pink arrow). Nevertheless, 1,6-hexanediol-elicited particles displayed irregular shapes and were more heterogeneous than those induced by diamide. Remarkably, at 8% (w/v), 1,6-hexanediol completely dissolved the vimentin network (Fig. 6D). The narrow range of concentrations in which these changes occur suggest a threshold effect of 1,6-hexanediol. In contrast, 2,5-hexanediol did not alter filament integrity at any of the concentrations used (Fig. 6D). Taken together, these results suggest that low complexity domains may be involved both in keeping filament assembly and in maintaining diamide-elicited vimentin dots, with the latter structures being sensitive to lower concentrations of 1,6-hexanediol.

**Figure 6.**
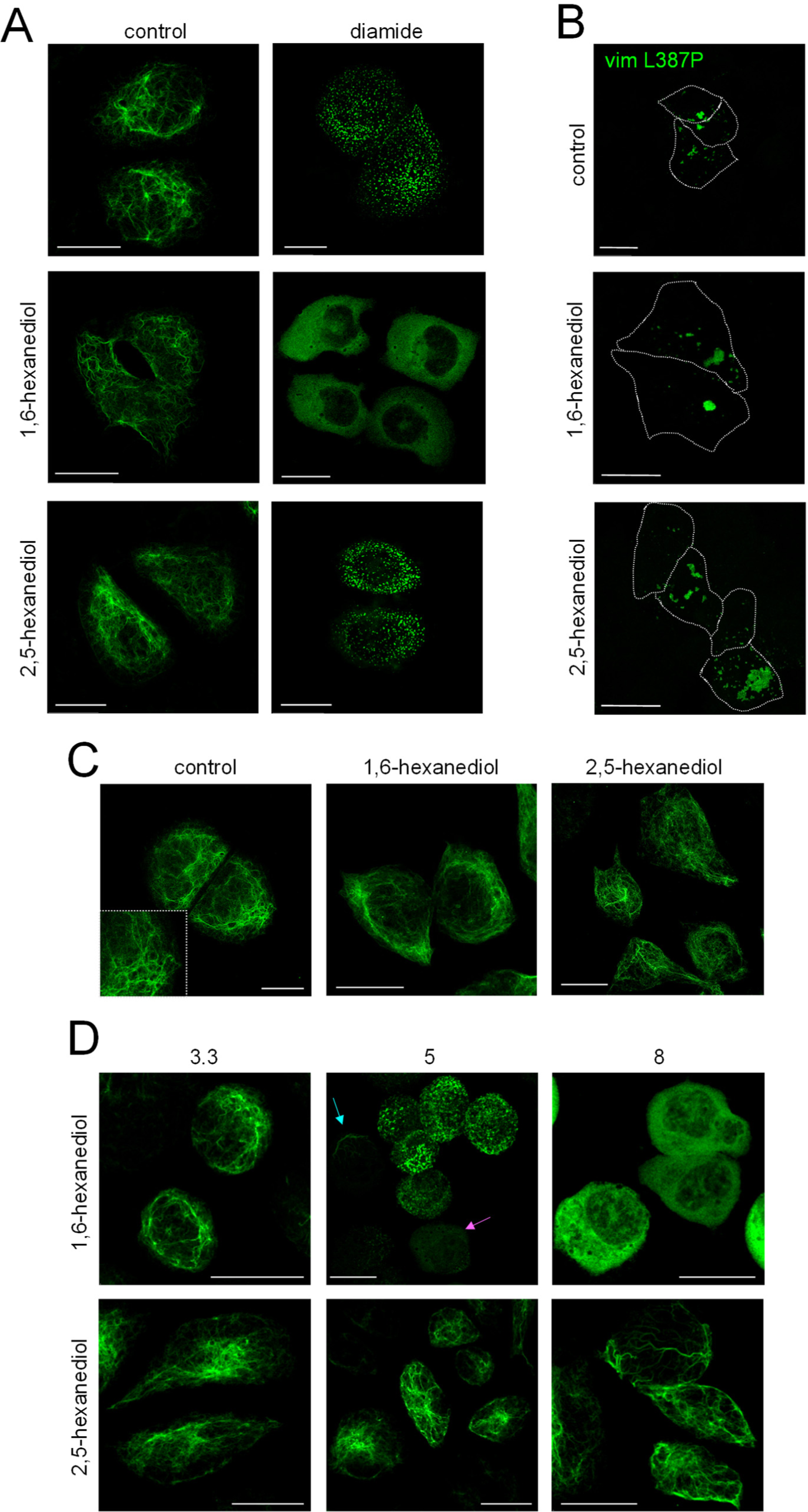
Effect of aliphatic alcohols on vimentin dots and vimentin filaments. (A) SW13/cl.2 cells stably expressing untagged vimentin wt were incubated for 20 min in the absence or presence 1 mM diamide. During the last five minutes of the incubation, 1,6-hexanediol or 2,5-hexanediol at 3.3% (w/v) final concentration, were added to the cell culture medium. (B) SW13/cl.2 cells transfected with a vimentin L387P mutant were incubated with the indicated alcohols as in (A). (C) and (D) Cells expressing vimentin wt were incubated with the indicated alcohols at 3.3% (w/v) final concentration for 20 min (C) or with the indicated concentrations (in %(w/v)) of the corresponding alcohols for 5 min (D). Immediately after treatments, cells were fixed and processed by immunofluorescence of vimentin detection by confocal microscopy. Blue arrow, cell with conserved filaments; pink arrow, cell with diffuse vimentin. Overall projections are shown. Bars, 20 µm.

### Implications of the presence of C328 in the formation of vimentin dots

We have previously shown that the single cysteine of vimentin is critical for its remodeling in response to various oxidants and electrophiles (18, 20, 21). Here we confirmed that the cysteine-deficient mutant vimentin C328A is basically resistant to diamide-elicited filament conversion into dots (Fig. 7A). Moreover, addition 3.3% (w/v) 1,6-hexanediol to diamide treated cells did not impair filament integrity, although a faint background was observed in some cells. Therefore, the vimentin C328A mutation not only protects filaments from diamide-elicited dot formation, but also from the subsequent dissolution by 3.3% (w/v) 1,6-hexanediol. This suggests that indeed, filament dissociation by diamide treatment facilitates 1,6-hexanediol action. Nevertheless, mature C328A vimentin filaments were susceptible to disruption by 1,6-hexanediol per se at higher concentrations (5 and 8% (w/v)), similar to wt (Fig. 7B). Thus, 5% (w/v) 1,6-hexanediol elicited the appearance of irregular particles or short filament fragments and 8% led to a diffuse vimentin C328A pattern (Fig. 7B). These observations indicate that C328 is not required for filament disruption by high concentrations of the aliphatic alcohol. In contrast, similar to the effect on vimentin wt, 2,5-hexanediol, did not alter vimentin C328A filaments under any experimental condition (Fig. 7A and B).

**Figure 7.**
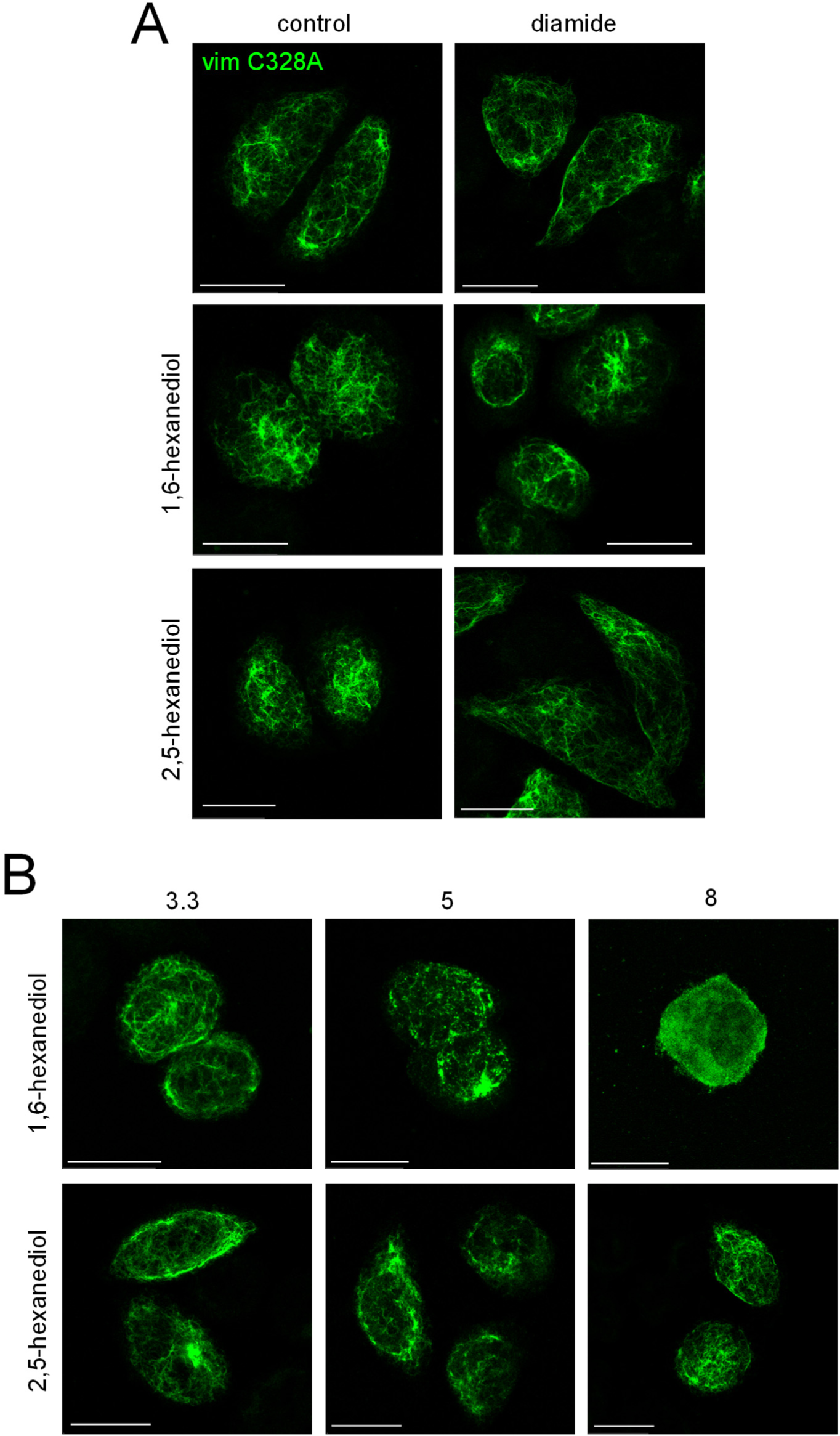
Effect of diamide and aliphatic alcohols on a vimentin C328A mutant. (A) SW13/cl.2 cells stably expressing vimentin C328A were incubated in the absence or presence of 1 mM diamide for 20 min. During the last five minutes of the incubation, 1,6-hexanediol or 2,5-hexanediol at 3.3% (w/v) final concentration, were added to the cell culture medium. (B) Cells expressing vimentin C328A were incubated with the indicated concentrations (in %(w/v)) of 1,6-hexanediol or 2,5-hexanediol for 5 min (D). After treatments, cells were fixed and processed by immunofluorescence of vimentin detection by confocal microscopy. Overall projections are shown. Bars, 20 µm.

## Discussion

Intermediate filaments are critical elements in the cellular response to stress. They form versatile arrangements that undergo fast remodeling into a variety of patterns, including bundles, cages, diffuse protein or extracellular structures, that exert protective effects through a variety of mechanisms (11, 28). Here we have observed that several oxidants induce the dissociation of vimentin filaments into small particles or dots, either preferentially at the cell periphery, as seen for H_2_O_2_, or throughout the cell, as in the case of diamide. Interestingly, diamide-elicited vimentin particles share the properties of biomolecular condensates because they are mostly spherical, they recover from photobleaching and appear to fuse. Importantly, diamide-elicited vimentin dots are reversible upon removal of the oxidant and are dispersed by certain aliphatic alcohols, suggesting the involvement of labile cross-ꞵ interactions in their formation (33). In view of these findings, we hypothesize that these droplet-like structures may represent a fast rearrangement of the vimentin network, preventing excessive damage to the structure of filament subunits.

We observed that the majority of diamide-elicited vimentin dots were virtually circular. Most dots displayed an area smaller than 0.1 µm^2^, and many of them were compatible with the size of a few ULF, indicating that filaments were dissociated into small segments. Although we do not know the precise structure of vimentin dots, we have previously reported that vimentin monomers appear to be intact in diamide-treated cells (26), thus supporting that the reorganization of filament into dots does not require protein cleavage. Interestingly, the effect of short term diamide treatment was reversible. Upon removal of the oxidant, dots elicited by a 5 min diamide treatment quickly reverted to squiggles and then formed filaments of normal appearance. Notably, when diamide treatment was continued for 15 min, recovery of the network was slower, and dots present, first coalesced forming bigger accumulations, which later appeared joined by irregular filaments. Nevertheless, this pattern clearly indicated a recovery since upon sustained incubation with diamide, vimentin dots evolved towards a diffuse cytoplasmic background, which could be the result of additional modifications. Notably, diamide can also disrupt actin filaments and microtubules (20, 26), the involvement of which in vimentin reorganization and filament recovery need further study.

Diamide-elicited vimentin dots are highly motile and dynamic structures. On one hand, some of them can travel long distances within cells, although at present it is not known whether these movements occur on remaining microtubule tracks or other structures. Vimentin precursors (ULF) are known to move along microtubules, whereas actin can restrict their transport (49). Therefore, it could be hypothesized that the movement of diamide-elicited vimentin dots would be the result of the modulation of the various cytoskeletal networks by oxidative stress. On the other hand, vimentin droplets are capable of FRAP. It is well known that vimentin filaments can exchange subunits along their length, and their dynamic behavior has been previously documented through FRAP assays (20, 46, 50, 51). In fact, in earlier works, we observed recovery of vimentin filaments from photobleaching in a matter of minutes (18, 20). Here, we observed that FRAP of diamide-elicited vimentin dots was very fast. In fact, an increase in fluorescence was obvious at the first time point recorded after bleaching (five seconds). These observations are consistent with studies reporting fast exchange of subunits after FRAP or photoconversion of particles or ULF formed by assembly-deficient vimentin mutants (52). Taken together, our results indicate that vimentin particles are highly dynamic, and likely contain “functional units”. Further support of the functionality of vimentin in the dots is provided by assays where these structures are exchanged between cells fused by PEG treatment. Fusion of cells expressing vimentin tagged with different “colors” allowed detection of particles bearing both tags, either as a result of particle coalescence or of exchange of subunits. Importantly, cell fusion strategies have been previously used to prove the ability of vimentin filaments from differentially tagged populations to severe and anneal end to end, resulting in filaments of alternate “colors”, as well as to propose an intercalary mode of subunit exchange (39). Nevertheless, to the best of our knowledge, this type of dynamic interactions of vimentin as a result of oxidative stress has not been previously studied.

Interestingly, the N-termini of vimentin monomers have been reported to project towards the interior of the filaments (36), where they appear to be hold together by transient interactions contributing to maintain filament assembly (35). Therefore, the associated N-termini could constitute a “phase-separated” compartment within the filament lumen. Interestingly, treatment with 1,6-hexanediol at 8% (w/v) has been reported to disassemble vimentin filaments in HeLa cells and fibroblasts (33, 35). This was interpreted as a result of the disruption of the polymer containing low complexity sequences. Our results show that under oxidative stress vimentin filaments reversibly remodel into particles, which retain properties of biomolecular condensates. Therefore, hypothetically, these particles could also be hold together by labile interactions between low complexity sequences. Importantly, interference with these labile interactions, either by posttranslational modifications of the head domains or by treatment with low concentrations of 1,6-hexanediol (3.3% (w/v)), could result in further particle disassembly, whereas reversal of oxidative stress would allow filament recovery. A scheme of this hypothesis in shown in Suppl. Fig. 1. In contrast, clearly higher concentrations of 1,6-hexanediol (5 to 8% (w/v)) are required to disrupt full filaments in cells. Therefore, diamide-elicited dots are more susceptible to the effect of 1,6-hexanediol than mature filaments. Nevertheless, the use of aliphatic alcohols as indicators of phase separation also has limitations (45), and their effects in cells need to be interpreted cautiously, since under certain conditions they can induce the formation of aggregates per se, inhibit kinases and/or phosphatases and elicit toxic effects (47, 53).

Importantly, redox alterations and protein oxidative modifications play a key role in the modulation of phase separation processes (54, 55). Conversely, the process of phase separation is involved in the maintenance of cellular redox status (56). Additionally, macromolecular crowding-driven phase separation (57) can have important consequences in the rate, extent and pathways of protein oxidation (58). Therefore, a complex reciprocal modulation of phase separation and redox regulation can take place. Our results could be viewed as a process of oxidation-dependent phase separation or partition of vimentin between different states, which can be modulated by influencing the cellular environment (18, 20, 26).

We have shown that the remodeling of vimentin filaments into particles with characteristics of condensates elicited by diamide requires the presence of the single cysteine of vimentin, C328, since the vimentin C328A mutant is resistant to this effect. This indicates that modifications of this residue may be involved in diamide-elicited vimentin remodeling. Phase separation can be modulated by a variety of posttranslational modifications (29), and in particular by modifications of thiol groups (54). Cysteine residues can undergo a plethora of structurally different modifications with non-overlapping functional consequences (59, 60). Regarding condensate formation, a recent work has reported that distinct thiol modifications can differentially impact the participation of proteins in phase separation (54). Importantly, C328 is a hot spot for posttranslational modifications and a key for vimentin remodeling in response to various oxidants and electrophiles, which may induce distinct vimentin network rearrangements (reviewed in (8, 61)). Accumulating evidence suggests that C328 modification by bulky or charge-modifying moieties could destabilize filaments or interfere with assembly, for instance by perturbing contacts between ULF during filament elongation, or hampering subunit exchange (18, 24, 61). In cells, C328 modifications can include sulfenylation, persulfidation, CoAlation or glutathionylation, among others (8, 61–64). Indeed, we have previously confirmed that treating cells with diamide elicits a significant oxidation of C328, as evidenced by the decrease in the proportion of vimentin monomers with a free C328 thiol group (26). Interestingly, vimentin assembly in cells appears to involve an important degree of overlap of consecutive ULF (40), for which C328 modification could provoke steric hindrance.

Remarkably, the observation that vimentin C328A filaments are not only resistant to diamide but also to diamide followed by 1,6-hexanediol treatment, supports that C328-dependent filament dissociation is required for disruption of vimentin structures by low concentrations of 1,6-hexanediol, which potentially could access the particle core more easily. Nevertheless, higher concentrations of the alcohol readily destabilize both vimentin wt and C328A filaments, suggesting that it acts through a mechanism not requiring redox regulation.

It should be considered that in cells, other posttranslational modifications could cooperate with C328 modifications in order to generate vimentin dots or to sensitize vimentin to disassembly, for instance, modifications taking place at the head domain (15, 22). In previous works we observed that vimentin filament reorganization by diamide was prevented by strategies inhibiting ATP synthesis, which also hampered subunit exchange (18). Therefore, ATP-dependent processes could participate in diamide-elicited vimentin remodeling. It is known that cellular condensates can be regulated by ATP-consuming processes, indicating that active steps are involved in their formation (reviewed in (29)). One of such ATP-dependent processes is protein phosphorylation. We previously reported that the phosphatase inhibitor calyculin A potentiated filament disruption by diamide, whereas a vimentin mutant in which eleven serine residues of the N-terminal of vimentin were substituted by alanine, was partially protected from diamide-elicited conversion into dots (18). These observations suggested that N-terminal phosphorylation of vimentin could contribute to diamide effect. Importantly, phosphorylation by PKA plays an important role in the disassembly of type III intermediate filaments in cells (22) as well as in vitro (34), and can inhibit the self-association of head domains (34). Therefore, phosphorylation of the vimentin head domains could affect low complexity domain interactions, as previously discussed (34).

The role of diamide-elicited vimentin dots, if any, is not completely understood at this point. Given the fact that these structures are reversible, this fast remodeling could represent a defense mechanism against severe oxidative damage of the protein, which could take place at C328 and/or other residues. Indeed, images of recovery after a 15 min diamide treatment show that vimentin dots reconnect and become linked by thin filaments. Therefore, it could be speculated that vimentin dots constitute seeding structures or reservoirs of functional protein, facilitating reformation of filaments after short term oxidative stress. On the other hand, under prolongued oxidative stress, as after long diamide treatment, vimentin could be more extensively disassembled. Therefore, a sequential disassembly of vimentin filaments could be envisaged (Suppl. Fig. 1). Certain oxidants eliciting C328 modifications, would first provoke a partial or total remodeling of vimentin filaments into particles, hypothetically still conserving a biomolecular condensate core, held together by low complexity domain interactions that would allow filament reassembly. Nevertheless, in these particles, N-terminal domains would be more accessible to further modifications or to molecules disrupting these low affinity interactions, facilitating the complete solubilization of the protein if stress persists.

## Acknowledgements.

We are indebted to Prof. G. Rivas and Dr. S. Zorrilla (CIB Margarita Salas, CSIC), for valuable comments and discussion. We thank Elena Sala Lara for preparation of the vimentin L387P mutant. Feedback from EpiLipidNet (CA19105) is gratefully acknowledged. We thank the personnel of the Optical Microscopy facility at Centro de Investigaciones Biológicas Margarita Salas for their advice.

## Abbreviations

FRAP: flurorescence recovery after photobleaching
GFP: green fluorescent protein
MBrB: monobromobimane
PEG: polyethylene glycol
ULF: unit length filament

## References

1. Liu Y, van den Ent F, Lowe J. Filament structure and subcellular organization of the bacterial intermediate filament-like protein crescentin. Proc Natl Acad Sci U S A. 2024;121(7):e2309984121. Doi: 10.1073/pnas.2309984121.

2. Eriksson JE, Dechat T, Grin B, Helfand B, Mendez M, Pallari HM, et al. Introducing intermediate filaments: from discovery to disease. J Clin Invest. 2009;119(7):1763–71. Doi: 10.1172/JCI38339.

3. Eldirany SA, Lomakin IB, Ho M, Bunick CG. Recent insight into intermediate filament structure. Curr Opin Cell Biol. 2021;68:132–43. Doi: 10.1016/j.ceb.2020.10.001.

4. Herrmann H, Aebi U. Intermediate filaments: molecular structure, assembly mechanism, and integration into functionally distinct intracellular Scaffolds. Annu Rev Biochem. 2004;73:749–89. Doi: 10.1146/annurev.biochem.73.011303.073823.

5. Szeverenyi I, Cassidy AJ, Chung CW, Lee BT, Common JE, Ogg SC, et al. The Human Intermediate Filament Database: comprehensive information on a gene family involved in many human diseases. Hum Mutat. 2008;29(3):351–60. Doi: 10.1002/humu.20652.

6. Toivola DM, Strnad P, Habtezion A, Omary MB. Intermediate filaments take the heat as stress proteins. Trends Cell Biol. 2010;20(2):79–91. Doi: 10.1016/j.tcb.2009.11.004.

7. Etienne-Manneville S. Cytoplasmic Intermediate Filaments in Cell Biology. Annu Rev Cell Dev Biol. 2018;34:1–28. Doi: 10.1146/annurev-cellbio-100617-062534.

8. Pérez-Sala D, Quinlan R. The redox-responsive roles of intermediate filaments in cellular stress detection, integration and mitigation. Curr Opin Cell Biol. 2024;86:102283. Doi: 10.1016/j.ceb.2023.102283.

9. Venturini A, Passalacqua M, Pelassa S, Pastorino F, Tedesco M, Cortese K, et al. Exosomes From Astrocyte Processes: Signaling to Neurons. Frontiers in Pharmacology. 2019;10:1452. Doi: 10.3389/fphar.2019.01452.

10. Ramos I, Stamatakis K, Oeste CL, Perez-Sala D. Vimentin as a Multifaceted Player and Potential Therapeutic Target in Viral Infections. International Journal of Molecular Sciences. 2020;21(13):4675. Doi: 10.3390/ijms21134675.

11. Parvanian S, Zha H, Su D, Xi L, Jiu Y, Chen H, et al. Exosomal Vimentin from Adipocyte Progenitors Protects Fibroblasts against Osmotic Stress and Inhibits Apoptosis to Enhance Wound Healing. International Journal of Molecular Sciences. 2021;22(9):4678. Doi: 10.3390/ijms22094678.

12. Lalioti V, González-Sanz S, Lois-Bermejo I, González-Jiménez P, Viedma-Poyatos A, Merino A, et al. Cell surface detection of vimentin, ACE2 and SARS-CoV-2 Spike proteins reveals selective colocalization at primary cilia. Scientific Reports. 2022;12:7063. Doi: 10.1038/s41598-022-11248-y.

13. Hol EM, Capetanaki Y. Type III Intermediate Filaments Desmin, Glial Fibrillary Acidic Protein (GFAP), Vimentin, and Peripherin. Cold Spring Harbor Perspectives in Biology. 2017;9(12):a021642. Doi: 10.1101/cshperspect.a021642.

14. Herrmann H, Aebi U. Intermediate Filaments: Structure and Assembly. Cold Spring Harbor perspectives in biology. 2016;8(11):a018242. Doi: 10.1101/cshperspect.a018242.

15. Sripathi SR, He W, Um JY, Moser T, Dehnbostel S, Kindt K, et al. Nitric oxide leads to cytoskeletal reorganization in the retinal pigment epithelium under oxidative stress. Adv Biosci Biotechnol. 2012;3:1167–78. Doi: 10.4236/abb.2012.38143.

16. Li J, Gao W, Zhang Y, Cheng F, Eriksson JE, Etienne-Manneville S, et al. Engagement of vimentin intermediate filaments in hypotonic stress. J Cell Biochem. 2019;120(8):13168–76. Doi: 10.1002/jcb.28591.

17. Zhang Y, Zhao S, Li Y, Feng F, Li M, Xue Y, et al. Host cytoskeletal vimentin serves as a structural organizer and an RNA-binding protein regulator to facilitate Zika viral replication. Proc Natl Acad Sci U S A. 2022;119(8). Doi: 10.1073/pnas.2113909119.

18. Mónico A, Duarte S, Pajares MA, Pérez-Sala D. Vimentin disruption by lipoxidation and electrophiles: role of the cysteine residue and filament dynamics. Redox Biol. 2019;23:101098. Doi: 10.1016/j.redox.2019.101098.

19. Kueper T, Grune T, Prahl S, Lenz H, Welge V, Biernoth T, et al. Vimentin is the specific target in skin glycation. Structural prerequisites, functional consequences, and role in skin aging. J Biol Chem. 2007;282(32):23427–36. Doi: 10.1074/jbc.M701586200.

20. Pérez-Sala D, Oeste CL, Martínez AE, Garzón B, Carrasco MJ, Cañada FJ. Vimentin filament organization and stress sensing depend on its single cysteine residue and zinc binding. Nature Commun. 2015;6:7287. Doi: 10.1038/ncomms8287.

21. González-Jiménez P, Duarte S, Martínez-Fernández A, Navarro-Carrasco E, Lalioti V, Pajares MA, et al. Vimentin single cysteine residue acts as a tunable sensor for network organization and as a key for actin remodeling in response to oxidants and electrophiles. Redox Biol. 2023;64:102756. Doi: 10.1016/j.redox.2023.102756.

22. Eriksson JE, He T, Trejo-Skalli AV, Härmälä-Braskén A-S, Hellman J, Chou Y-H, et al. Specific in vivo phosphorylation sites determine the assembly dynamics of vimentin intermediate filaments. J Cell Sci. 2004;117:919–32. Doi:

23. Chou YH, Khuon S, Herrmann H, Goldman RD. Nestin promotes the phosphorylation-dependent disassembly of vimentin intermediate filaments during mitosis. Mol Biol Cell. 2003;14(4):1468–78. Doi: 10.1091/mbc.E02-08-0545.

24. Kaus-Drobek M, Mucke N, Szczepanowski RH, Wedig T, Czarnocki-Cieciura M, Polakowska M, et al. Vimentin S-glutathionylation at Cys328 inhibits filament elongation and induces severing of mature filaments in vitro. FEBS J. 2020;287:5304–22. Doi: 10.1111/febs.15321.

25. Lois-Bermejo I, González-Jiménez P, Duarte S, Pajares MA, Pérez-Sala D. Vimentin tail segments are differentially exposed at distinct cellular locations and in response to stress. Frontiers Cell Dev Biology. 2022;10:908263. Doi: 10.3389/fcell.2022.908263.

26. Martínez AE, González-Jiménez P, Vidal-Verdú C, Pajares MA, Pérez-Sala D. Intracellular pH modulates vimentin remodeling in response to oxidants. BioRxiv. 2023. Doi: 10.1101/2023.12.21.572888.

27. Quinlan R. Cytoskeletal competence requires protein chaperones. Prog Mol Subcell Biol. 2002;28:219–33. Doi: 10.1007/978-3-642-56348-5_12.

28. Pattabiraman S, Azad GK, Amen T, Brielle S, Park JE, Sze SK, et al. Vimentin protects differentiating stem cells from stress. Scientific Reports. 2020;10(1):19525. Doi: 10.1038/s41598-020-76076-4.

29. Alberti S, Dormann D. Liquid-Liquid Phase Separation in Disease. Annu Rev Genet. 2019;53:171–94. Doi: 10.1146/annurev-genet-112618-043527.

30. Kanaan NM, Hamel C, Grabinski T, Combs B. Liquid-liquid phase separation induces pathogenic tau conformations in vitro. Nat Commun. 2020;11(1):2809. Doi: 10.1038/s41467-020-16580-3.

31. Woodruff JB, Ferreira Gomes B, Widlund PO, Mahamid J, Honigmann A, Hyman AA. The Centrosome Is a Selective Condensate that Nucleates Microtubules by Concentrating Tubulin. Cell. 2017;169(6):1066–77 e10. Doi: 10.1016/j.cell.2017.05.028.

32. Mitrea DM, Mittasch M, Gomes BF, Klein IA, Murcko MA. Modulating biomolecular condensates: a novel approach to drug discovery. Nat Rev Drug Discov. 2022;21(11):841–62. Doi: 10.1038/s41573-022-00505-4.

33. Lin Y, Mori E, Kato M, Xiang S, Wu L, Kwon I, et al. Toxic PR Poly-Dipeptides Encoded by the C9orf72 Repeat Expansion Target LC Domain Polymers. Cell. 2016;167(3):789–802 e12. Doi: 10.1016/j.cell.2016.10.003.

34. Zhou X, Lin Y, Kato M, Mori E, Liszczak G, Sutherland L, et al. Transiently structured head domains control intermediate filament assembly. Proc Natl Acad Sci U S A. 2021;118(8). Doi: 10.1073/pnas.2022121118.

35. Zhou X, Kato M, McKnight SL. How do disordered head domains assist in the assembly of intermediate filaments? Curr Opin Cell Biol. 2023;85:102262. Doi: 10.1016/j.ceb.2023.102262.

37. Swiader A, Camare C, Guerby P, Salvayre R, Negre-Salvayre A. 4-Hydroxynonenal Contributes to Fibroblast Senescence in Skin Photoaging Evoked by UV-A Radiation. Antioxidants (Basel). 2021;10(3):365. Doi: 10.3390/antiox10030365.

38. Griesser E, Vemula V, Mónico A, Pérez-Sala D, Fedorova M. Dynamic posttranslational modifications of cytoskeletal proteins unveil hot spots under nitroxidative stress. Redox Biol. 2021;44:102014. Doi: 10.1016/j.redox.2021.102014.

39. Colakoglu G, Brown A. Intermediate filaments exchange subunits along their length and elongate by end-to-end annealing. J Cell Biol. 2009;185(5):769–77. Doi: 10.1083/jcb.200809166.

40. Nunes Vicente F, Lelek M, Tinevez JY, Tran QD, Pehau-Arnaudet G, Zimmer C, et al. Molecular organization and mechanics of single vimentin filaments revealed by super-resolution imaging. Sci Adv. 2022;8(8):eabm2696. Doi: 10.1126/sciadv.abm2696.

41. Terriac E, Coceano G, Mavajian Z, Hageman TA, Christ AF, Testa I, et al. Vimentin Levels and Serine 71 Phosphorylation in the Control of Cell-Matrix Adhesions, Migration Speed, and Shape of Transformed Human Fibroblasts. Cells. 2017;6(1):2. Doi: 10.3390/cells6010002.

42. Kosower NS, Kosower EM, Wertheim B, Correa WS. Diamide, a new reagent for the intracellular oxidation of glutathione to the disulfide. Biochem Biophys Res Commun. 1969;37(4):593–6.

43. Kosower NS, Kosower EM. Diamide: an oxidant probe for thiols. Methods Enzymol. 1995;251:123–33.

44. Irgen-Gioro S, Yoshida S, Walling V, Chong S. Fixation can change the appearance of phase separation in living cells. eLife. 2022;11. Doi: 10.7554/eLife.79903.

45. Alberti S, Gladfelter A, Mittag T. Considerations and Challenges in Studying Liquid-Liquid Phase Separation and Biomolecular Condensates. Cell. 2019;176(3):419–34. Doi: 10.1016/j.cell.2018.12.035.

46. Yoon M, Moir RD, Prahlad V, Goldman R. Motile properties of vimentin intermediate filament networks in living cells. J Cell Biol. 1998;143:147–57.

47. Duster R, Kaltheuner IH, Schmitz M, Geyer M. 1,6-Hexanediol, commonly used to dissolve liquid-liquid phase separated condensates, directly impairs kinase and phosphatase activities. J Biol Chem. 2021;296:100260. Doi: 10.1016/j.jbc.2021.100260.

48. Cogne B, Bouameur JE, Hayot G, Latypova X, Pattabiraman S, Caillaud A, et al. A dominant vimentin variant causes a rare syndrome with premature aging. Eur J Hum Genet: EJHG. 2020;28:1218–30. Doi: 10.1038/s41431-020-0583-2.

49. Robert A, Herrmann H, Davidson MW, Gelfand VI. Microtubule-dependent transport of vimentin filament precursors is regulated by actin and by the concerted action of Rho- and p21-activated kinases. FASEB J. 2014;28(7):2879–90. Doi: 10.1096/fj.14-250019.

50. Helfand BT, Mendez MG, Murthy SN, Shumaker DK, Grin B, Mahammad S, et al. Vimentin organization modulates the formation of lamellipodia. Mol Biol Cell. 2011;22(8):1274–89. Doi: 10.1091/mbc.E10-08-0699.

51. Yang CY, Chang PW, Hsu WH, Chang HC, Chen CL, Lai CC, et al. Src and SHP2 coordinately regulate the dynamics and organization of vimentin filaments during cell migration. Oncogene. 2019;38(21):4075–94. Doi: 10.1038/s41388-019-0705-x.

52. Robert A, Rossow MJ, Hookway C, Adam SA, Gelfand VI. Vimentin filament precursors exchange subunits in an ATP-dependent manner. Proc Natl Acad Sci U S A. 2015;112(27):E3505–14. Doi: 10.1073/pnas.1505303112.

53. Meduri R, Rubio LS, Mohajan S, Gross DS. Phase-separation antagonists potently inhibit transcription and broadly increase nucleosome density. J Biol Chem. 2022;298(10):102365. Doi: 10.1016/j.jbc.2022.102365.

54. Vignane T, Hugo M, Hoffmann C, Katsouda A, Petric J, Wang H, et al. Protein thiol alterations drive aberrant phase separation in aging. bioRxiv. 2023. Doi: 10.1101/2023.11.07.566021.

55. Fuentes-Lemus E, Davies MJ. Effect of crowding, compartmentalization and nanodomains on protein modification and redox signaling - current state and future challenges. Free Radic Biol Med. 2023;196:81–92. Doi: 10.1016/j.freeradbiomed.2023.01.011.

56. Saito Y, Kimura W. Roles of Phase Separation for Cellular Redox Maintenance. Frontiers in genetics. 2021;12:691946. Doi: 10.3389/fgene.2021.691946.

57. Monterroso B, Margolin W, Boersma AJ, Rivas G, Poolman B, Zorrilla S. Macromolecular Crowding, Phase Separation, and Homeostasis in the Orchestration of Bacterial Cellular Functions. Chem Rev. 2024;124(4):1899–949. Doi: 10.1021/acs.chemrev.3c00622.

58. Fuentes-Lemus E, Reyes JS, Gamon LF, Lopez-Alarcon C, Davies MJ. Effect of macromolecular crowding on protein oxidation: Consequences on the rate, extent and oxidation pathways. Redox Biol. 2021;48:102202. Doi: 10.1016/j.redox.2021.102202.

59. Srivastava SK, Ramana KV, Chandra D, Srivastava S, Bhatnagar A. Regulation of aldose reductase and the polyol pathway activity by nitric oxide. Chem Biol Interact. 2003;143–144:333–40.

60. Oeste CL, Pérez-Sala D. Modification of cysteine residues by cyclopentenone prostaglandins: interplay with redox regulation of protein function. Mass Spectrom Rev. 2014;33:110–25. Doi: 10.1002/mas.21383.

61. Viedma-Poyatos A, Pajares MA, Pérez-Sala D. Type III intermediate filaments as targets and effectors of electrophiles and oxidants. Redox Biol. 2020;36:101582. Doi: 10.1016/j.redox.2020.101582.

62. Pedre B, Talwar D, Barayeu U, Schilling D, Luzarowski M, Sokolowski M, et al. 3-Mercaptopyruvate sulfur transferase is a protein persulfidase. Nat Chem Biol. 2023;19(4):507–17. Doi: 10.1038/s41589-022-01244-8.

63. Tossounian MA, Baczynska M, Dalton W, Newell C, Ma Y, Das S, et al. Profiling the Site of Protein CoAlation and Coenzyme A Stabilization Interactions. Antioxidants (Basel). 2022;11(7):1362. Doi: 10.3390/antiox11071362.

64. Fratelli M, Demol H, Puype M, Casagrande S, Eberini I, Salmona M, et al. Identification by redox proteomics of glutathionylated proteins in oxidatively stressed human T lymphocytes. Proc Natl Acad Sci U S A. 2002;99:3505–10.

