## Supplementary material for "Formation of vimentin biomolecular condensate-like structures under oxidative stress": Suppl. Fig.

Centro de Investigaciones Biológicas Margarita Salas, CSIC, 28040 Madrid, Spain

Running title: Vimentin condensates in oxidative stress

**List of Supplementary Videos:**

**Supplementary Video 1. Time lapse of diamide-elicited vimentin filament dissociation.** Live SW13/Cl.2 cells transiently transfected with RFP//vimentin (coding for untagged vimentin) plus GFP-vimentin were treated with 1 mM diamide. One minute later, images of single sections were acquired every 10 s in a SP5 microscope equipped with a thermostatized chamber. Images (3 frames per s) were converted to video with Image J.

**Supplementary Video 2. Time lapse of particle movement in diamide-treated cells.** Cells transiently transfected with RFP//vimentin plus GFP-vimentin and treated with diamide were monitored live. Images of single sections were acquired every 5 s in a SP5 microscope equipped with a thermostatized chamber. Images (3 frames per s) were converted to video with Image J.

**Supplementary Video 3. Time lapse of particle movement in diamide-treated cells.** Cells transiently transfected with RFP//vimentin plus GFP-vimentin and treated with diamide were monitored live. Images of single sections were acquired every 10 s in a SP5 microscope equipped with a thermostatized chamber. Images (10 frames per s) were converted to video with Image J.

**Supplementary Video 4. Tracking of diamide elicited vimentin particles.** Live cells expressing untagged and GFP-tagged vimentin were monitored after treatment with diamide. Single sections were acquired every 5 s. Movement of particles was analyzed with the Particle Trackmate plugin of Image J. Trajectories are colored according to their length as specified in Fig. 3, with those approaching 10 µm appearing in red. Movement of particles is denoted by circles. Images (5 frames per s) were converted to video with Image J.

**Supplementary Video 5. FRAP assay of diamide elicited vimentin particles.** Live cells expressing a combination of untagged and GFP-tagged vimentin and treated with diamide were subjected to fluorescence bleaching as described in Fig. 4. After bleaching, single sections were acquired every 5 s. The video was prepared with Image J at 3 frames per second.

Supplementary Information

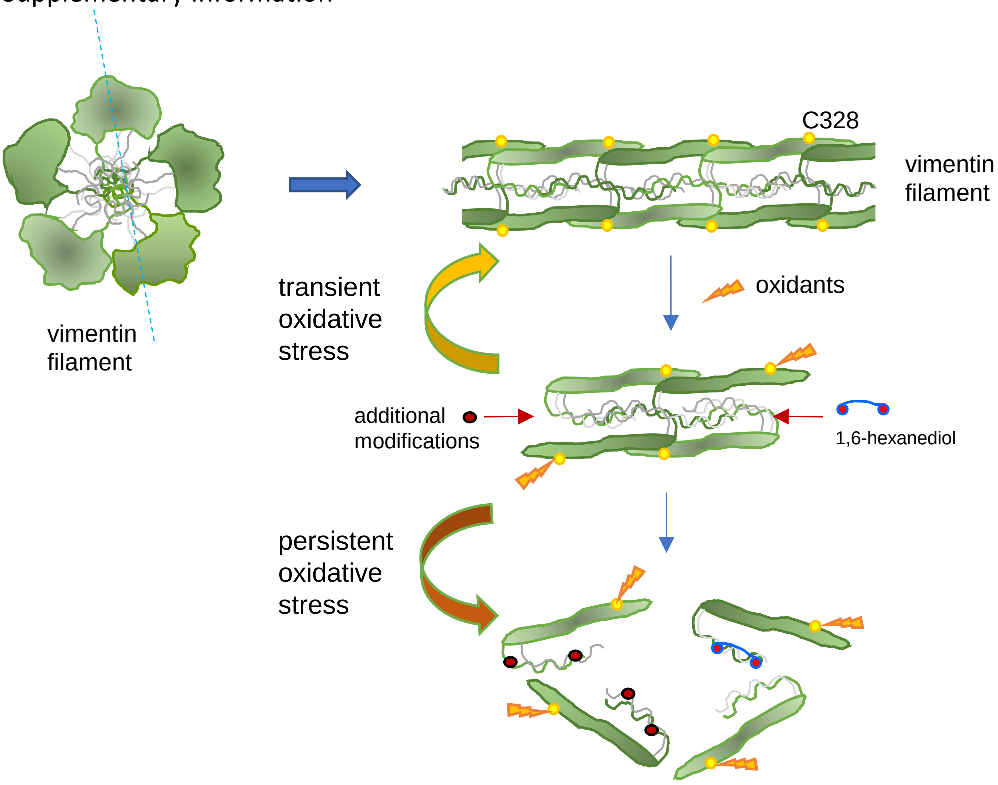


**Supplementary Figure 1. Scheme of the hypothetical reorganization of vimentin filaments in response to diamide.** The cartoon at upper left represents the transversal view of a vimentin filament, inspired by the recent report by Eibauer et al. (1), consisting of five "protofibrils", each of them formed in turn by two tetramers. The N-termini of vimentin monomers are projected towards the lumen of the filament forming a mesh bound by labile interactions that contribute to stabilize filament assembly (2). A cartoon representing a longitudinal view of the filament opened along the blue dotted line is shown at the right. The green aligned shapes represent the protofibrils, with the N-termini projecting towards the interior of the filament, and cysteine residues or clusters, exposed at the surface of the filament, represented in yellow. Please note that this representation does not intend to be structurally accurate. Oxidants, such as diamide, eliciting a significant modification of cysteine residues, could destabilize the filament leading to its dissociation into short particles with the characteristics of biomolecular condensates. In these particles, N-termini could still interact to a certain degree, holding the particle together. Under conditions of transient oxidative stress, particles would keep the capacity to reassemble upon removal of the oxidant, forming full filaments. Nevertheless, upon treatment with the aliphatic alcohol 1,6-hexanediol, or after further modifications such as phosphorylations or additional oxidations under persistent oxidative stress, particles would dissociate leading to diffuse vimentin.

**References**

1. Eibauer M, Weber MS, Kronenberg-Tenga R, Beales CT, Boujemaa-Paterski R, Turgay Y, et al. Vimentin filaments integrate low-complexity domains in a complex helical structure. Nat Struct Mol Biol. 2024. Doi: 10.1038/s41594-024-01261-2.
2. Zhou X, Kato M, McKnight SL. How do disordered head domains assist in the assembly of intermediate filaments? Curr Opin Cell Biol. 2023;85:102262. Doi: 10.1016/j.ceb.2023.102262.
